# Dietary inclusion of *Asparagopsis taxiformis* significantly reduces methane emissions in dairy ruminants by mechanistically altering vitamin B12-dependent and other methanogenesis precursor pathways

**DOI:** 10.1101/2025.05.29.656281

**Authors:** Katie Lawther, Nicholas J Dimonaco, Abdulai Guinguina, Sophie J Krizsan, Sharon A Huws

## Abstract

Ruminant products are consumed widely on a global level due to their high protein and micronutrient density. However, ruminant production is a major source of greenhouse gas emissions, particularly with respect to methane (CH₄), with ruminants contributing 33% of all anthropogenic CH₄ emissions. CH₄ is produced due to the natural fermentative processes undertaken by the complex rumen microbiome, primarily via the utilisation of hydrogen by rumen archaea to form CH₄. Previous studies have shown that feeding the red seaweed *Asparagopsis taxiformis* (ASP) to ruminants can reduce CH₄emissions from beef cattle by up to 80% (Roque et al., 2021). Nevertheless, the mechanism of action of this seaweed in terms of effects on the rumen microbiome is largely unknown, which is the main focus of this study. Six Nordic Red cows at 122 ± 13.7 (mean ± SD) days in milk were divided into 3 blocks by milk yield in Latin square design and fed grass silage and a commercial concentrate (60:40) either with or without 0.5% ASP on an organic matter basis. Rumen fluid was collected 19 days into each experimental period, with a holistic approach using both assembly and read mapping based approaches to interrogate taxonomic, functional, and ecological shifts applied to metagenomic data. We show that ASP reduces methane production not only through direct inhibition of methanogens but also by disrupting cobamide-dependent metabolic pathways and redirecting carbon flow toward pyruvate and propionate rather than acetate and methane. For the first time, specific enzymes involved in vitamin B12 (cobamide) biosynthesis are identified as suppressed by ASP, and microbial taxa contributing to these functional changes are elucidated showing both niche displacement and resilience within the rumen microbiome. These findings offer new mechanistic insight into how red seaweed supplementation modulates the rumen microbiome, supporting its potential role in sustainable ruminant production.

## Introduction

It is estimated that over 9.1 percent of humans do not have access to sufficient food and hence suffer from nutrient deficiencies and conditions such as anaemia and stunting (FAO, 2024). Moreover, it is predicted that the world’s population will reach 10.4 billion in the 2080s (United Nations Department of Economic and Social Affairs, Population Division, 2022), putting a further pressure on resources. Concomitantly, food production contributes up to 30% of global greenhouse gas (GHG) emissions (Clark et al., 2020), with agriculture playing a major role, mainly through livestock production and manure or slurry storage practices.

Ruminant products are consumed widely on a global level due to their high protein and micronutrient density, however, ruminant production is a major source of enteric methane (CH_4_), a greenhouse gas (GHG) that has between 27 and 30 times the global warming potential (GWP) of carbon dioxide (CO_2_), albeit it is a short-lived gas with a lifespan of approx. 12 years (Lynch et al., 2020). Indeed, ruminants account for 33% of all anthropogenic CH_4_ sources (Jackson et al., 2020; Crippa et al., 2021). However, when CH_4_ decomposes to CO_2_ in the atmosphere, it will bind to vegetation and will not contribute to further warming, suggesting that reducing CH_4_ emissions will have an immediate effect on global warming (Sterner and Johansson, 2017). Consequently, achieving food security and planetary health to limit malnutrition, whilst fulfilling government and global climate legislation, such as the Paris agreement which aims to limit global warming to 1.5°C above the pre-industrial baseline, is a major humanitarian challenge.

Ruminants are unique animals, as they possess a forestomach composed of 4 compartments: the reticulum, the rumen, the omasum and the abomasum. The rumen is the main fermentative compartment of the forestomach, composed of a complex, dynamic ecosystem of anaerobic bacteria, protozoa, anaerobic fungi, methanogenic archaea and bacteriophages. These microbes interact with each other and establish a symbiotic relationship with the host, providing it with energy in the form of volatile fatty acids arising from the breakdown of complex carbohydrates within dietary plant material. In this process hydrogen is released and utilised by methanogenic archaea to produce CH_4_, but many publications have shown that it is possible to reduce this biochemical process without determinately affecting the health of the animal. For example, efficient reductions in enteric emissions of CH_4_ from ruminants have been achieved through innovative feeding strategies, such as addition of the chemical 3- nitroxypropanol (Bovaer®) to the diet, resulting in approximately 25-30% reductions in CH_4,_ irrespective of the units used to express CH_4_ emissions (Hristov et al., 2015; Ardnt et al., 2021). More recently, studies have shown that feeding macroalgae, particularly the tropical macroalga *Asparagopsis taxiformis,* can reduce methane emissions from ruminants by between a 55% and 65% reduction *in vivo* when added to the diet of dairy cows (Stefenoni et al., 2021), and at most a 98% reduction *in vivo* when added to the diet of steers, again irrespective of expression units (Roque et al., 2021), all at an inclusion level of 0.5% of organic matter (OM).

To develop an anti-methanogenic dietary intervention, it is imperative that the mechanism of action is known and that there are no detrimental trade-offs to the fermentative capacity of the rumen microbiome and to animal health and productivity as a whole (Belanche et al., 2025). In terms of assessing mechanism of action of *A.taxiformis* on the rumen microbiome, few studies exist and those which are in public domain have utilised mainly metataxonomy only to monitor effects of bacterial and archaeal diversity. For example, using the *in vitro* RUSITEC technology which mimics the rumen environment, coupled with metataxonomy, Roque et al., (2019) showed that *A. taxiformis* did not substantially affect the rumen microbial diversity, nonetheless O’Hara et al., (2023) found a total reduction in all methanogenic archaea with increases in *Prevotella*, *Bifidobacterium*, *Succinivibrio*, *Ruminobacter* and unclassified *Lachnospiraceae*. Using *in vivo* studies, Krizsan et al. (2023) noted that rumen *Prevotella* were reduced in abundance and that there was a switch from a *Methanobrevibacter* dominated microbiome to that dominated by *Methanomethylophilaceae*, post feeding lactating dairy cows 0.5% OM of *A. taxiformis*. Conversely, using metataxonomic approaches, Indugu et al. (2024), found that feeding *A. taxiformis* again at 0.5% DM to lactating dairy cows reduced *Methanoshaeara* densities. Using shotgun metagenomics Indugu et al. (2024) also showed that *A. taxiformis* feeding reduced the copy number of Methyl reductase (MCR) gene which encodes the enzyme responsible for the final step in CH_4_ production by catalyzing the incorporation of coenzyme M and coenzyme B, resulting in heterosulfide production and release of CH_4_. Therefore, whilst some data exists on the mechanism of action of *A. taxiformis*, these are very contradictory as a whole and require advanced technologies to be applied in order to understand the fundamental basis on which *A. taxiformis* exudes an effect on the rumen microbiome, resulting in significant reduction in CH_4_ emissions.

Consequently, this study aimed to assess the effects of feeding *A. taxiformis* at 0.5% OM inclusion level on the rumen microbiome using shotgun metagenomic, thereby providing more certainty on the actual mechanism of action.

## Materials and methods

### Study design

This study complements the study performed by Krizsan et al. (2023) and provides novel mechanistic information on the mode of action of *A. taxiformis* in the rumen. As such more in depth information on the animal parameters can be found in Krizsan et al. (2023). In brief, the animal experiment was conducted at Röbäcksdalen experimental farm of the Swedish University of Agricultural Sciences in Umeå (63°45′N, 20°17′E) utilising six Nordic Red dairy cows weighing (mean ± SD) 611 kg ± 62.1 kg. The cows were 122 ± 13.7 days in milk, parity 2.7 ± 0.52, and producing 36.3 kg ± 2.47 kg of milk per day pretrial, and blocked experimentally by milk yield (MY). The cows were kept in an insulated free-stall barn, offered a total mixed ration (TMR) ad libitum, mixed with a feed mixer (Nolan A/S, Viborg, Denmark). The cows had free access to drinking water and were milked twice per day, at 06:00 and 16:30, throughout the experiment. The cows were randomly allocated to a dietary treatment within block, i.e., square, and assigned to an extra-period Latin-square change-over design consisting of three equal squares. The experimental periods lasted 21 days, with the last 7 days used for data recording and sampling. Rumen fluid was collected 19 days into each experimental period in triplicate (P1, P2, P3). All cows were fed a control diet composed of 600 g/kg dry matter (DM) of grass silage, 390 g/kg DM of a commercial concentrate (Komplett Amin 180; Lantmännen Lantbruk AB, Malmö, Sweden), and 10 g/kg DM of mineral mix (Mixa Optimal; Lantmännen Lantbruk AB, Malmö, Sweden). Dietary treatments comprised either no supplementation (control, CNT) or supplementation with the macroalga *A. taxiformis* (ASP) at 0.5% of organic matter (OM) intake.

### Rumen sampling

Rumen fluid was collected from all cows on day 19 in each period. The collection and preparation of rumen fluid was carried out in accordance with Chagas et al. (2021). Essentially rumen fluid samples were collected using a stomach tube (RUMINATOR; Munich, Germany) with the first sample of rumen fluid, comprising about 500 mL discarded, in order to avoid saliva contamination. Then, a sample of 500 mL was taken and filtered through a two-layer cheesecloth. Subsamples for microbial analysis were transferred to 2.0 mL Eppendorf tubes, immediately frozen using liquid Nitrogen, and kept at −80 °C in a freezer until analysis.

### Rumen Microbiome Sequencing

DNA was extracted using the Qiagen DNeasy PowerSoil Pro Kit, following the manufacturer’s guidelines. Briefly, 500 µL of thawed rumen fluid was centrifuged at 15,000 × g for 5 minutes to pellet the material, after which the supernatant was discarded. The pellet was then resuspended in 800 µL of CD1 solution, vortexed briefly, and transferred to a PowerBead Pro tube. Mechanical cell disruption and homogenization were performed through bead-beating at 5.5 m/s for three 1-minute cycles in a FastPrep instrument (MP Biomedicals), with ice incubation between cycles. A 75 µL aliquot of ZymoBIOMICs Microbial Community Standard (D6300) was used as a positive extraction control, while a reagent-only control “kitome” was also included, both controls were included in all downstream sequencing and analysis.

DNA concentration was measured using a Nanodrop spectrophotometer (ThermoFisher Scientific, NanoDrop™ One/OneC Microvolume UV-Vis Spectrophotometer, ND-ONE-W), and samples were diluted to a concentration of 8 ng/µL in TRIS 10 mM pH 8.0 where required, before being sent to the Queen’s University (QUB) Genomics Core Technology Unit (GCTU) for metagenomic sequencing. Thirty-eight samples underwent sequencing, for each of the 2 treatments 3 animals at 3 time points (technical replicates of 2) and a positive and negative control. The DNA underwent library preparation and shotgun metagenomic sequencing at the GCTU using an Illumina NovaSeq 6000 S4 300, (Paired end sequencing was performed, with a read length of 150 base pairs).

### Rumen Microbiome Sequence Analysis

DNA was extracted from a total of 36 rumen fluid samples and was sequenced producing a total of 2,084,236,576 raw reads, with an average of 57,895,460 reads per sample. FastQC was used to generate an overall sequence quality and distribution report which showed the reads had on average a PHRED score of ∼Q36. Next, Trimmomatic (v0.39) with the following parameters: ILLUMINACLIP: TruSeq3-PE-2:2:30:10 LEADING:3 TRAILING:3 SLIDINGWINDOW:4:20 MINLEN:50 was used to trim low-quality bases and adapters and identify reads with corresponding pairs (dropping reads if one pair was missing during sequencing). The reads were trimmed with a ‘SLIDINGWINDOW’ if at any site of 4 sequential bases the average quality dropped below Q20. Any reads reporting a quality score of < Q20 and short reads less than 50bp (after trimming) were discarded.

The ‘cleaned’ reads from each of the samples were then mapped to the human (GCF_000001405.40_GRCh38.p14), cow (GCF_002263795.2_ARS-UCD1.3) and sheep (GCF_016772045.1_ARS-UI_Ramb_v2.0) reference genomes using Bowtie2 (v2.5.1) with default parameters to identify the level of contamination and remove human and host-associated DNA. Reads that aligned with these reference genomes were considered contaminants and discarded. This resulted in a total of 1,956,850,528 clean reads, an average of 54,356,959 reads from each sample. These quality-checked, and decontaminated reads were metagenomically assembled using metaSPAdes (spades v4.0.6). This resulted in 786,040 contigs with an average of ∼21,834 per sample that were greater or equal to 2,500bp in length. The contigs underwent taxonomic classification using Kraken2 (v2.1.3) using the ‘k2_pluspfp_20240904’ (Standard plus Refeq protozoa, fungi and plant) precomputed database with default parameters.

Additionally, Pyrodigal (v3.0.1) was used to predict protein coding genes from the same contigs which were subsequently annotated with eggNOG-mapper (v2.1.12) using the eggNOG 5.0 database (Huerta-Cepas et al., 2019). Finally, Bowtie2 was applied to map the cleaned reads back to the assembled contigs to count the number of reads assigned to each taxa that had been identified by Kraken2. Next, bedtools intersect (v2.31.1, Quinlan and Hall, 2010) was used to identify reads that specifically mapped to protein-coding regions previously predicted by Prodigal on the assembled contigs. This step ensured that gene-level abundance estimates were accurately captured regardless of differences in contig assembly. The MetaPont package (pypi.org/project/MetaPont/) was developed to process the outputs of this workflow and includes a suite of tools for merging and analysing results from the various steps, producing tab-separated value (TSV) files for downstream interpretation. These files contained the full taxonomic and functional classifications alongside read mapping counts for each sample. A systematic extraction of the taxonomic contributors to functions of interest was then performed, providing insights into which taxa were driving key functional traits in each sample.

### Methanogen genome read mapping

To investigate the presence and species-level diversity of known rumen methanogens, organisms that are often low in abundance and challenging to reconstruct through metagenomic assembly, we performed read mapping against a curated set of 27 rumen-associated methanogen genomes (Leahy et al., 2010). Reads were aligned using Bowtie2 with the --very-sensitive-local parameter, allowing for greater sequence divergence between reads and reference genomes. The number of reads mapping to each contig within each genome was recorded. Subsequently, bedtools genomecov was used to calculate coverage statistics, generating a genome-wide coverage value that accounted for all contigs assigned to each methanogen genome. The ratio of total number of mapped reads (to the genome) to the total number of reads per sample (MP:TR) was calculated for each genome.

The complete metagenomic workflow is available at: github.com/TheHuwsLab/Metagenomic_Workflow.

### Statistical analysis and visualisation

The negative control was investigated for contamination and the taxonomy assigned to the positive control was compared to the expected taxonomic distribution for the ZymoBIOMICs Microbial Community Standard (D6300). All analysis was performed in R studio (v2024.9.1.394) (R version 4.4.2 (2024-10-31)) and visualised using ggplot2 (v3.5.1) unless otherwise specified. To consider biases, including library size difference, samples (reads described above) underwent normalisation using the trimmed mean of M-values (Pereira *et al*., 2018; Tapio *et al.,* 2023) using the edgeR package (v4.4.1) using the ‘calcNormFactors’ function. Following normalisation bacterial and archaeal data were separated, and phyla, family and genus levels were selected for further analysis. Relative abundance was calculated for each domain at each level, and visualised in stack bar charts where needed to add visualisation the top 20 taxa were plotted and the remaining were grouped together as “Other”. Alpha diversity indices including chao1, inverse Simpson and Shannon were calculated using the vegan package (v2.6.8) and visualised in box plots. The read data was then transformed using log(. + 1) for downstream statical analysis and principal component analysis (PCA) was performed using prcomp in Rstudio. Beta-diversity was performed with principal coordinate analysis (PCoA) based on Bray–Curtis dissimilarity matrices also using the vegan package (v2.6.8). The effect of treatment and time was assessed using Permutational multivariate analysis of variance (PERMANOVA) and was conducted using adonis() function in the vegan (v2.6.8) package, employing 999 iterations. Pairwise comparison were completed using the pairwiseAdonis (v0.4.1) package, and p values adjusted using the Benjamini-Hochberg method. Both Linear discriminant analysis Effect Size (LEfSe) analysis was completed and LDA plots visualised using the python scripts: lefse_format_input.py, lefse_run.py, lefse_plot_res.py, lefse_plot_features.py, using the lefse conda (v25.1.0) package ([v1.1.2], Segata *et al*., 2011, <u>SegataLab/lefse</u>). Default parameters were utilised and the option -o 1000000 included as recommended in by the authors manual, this scales the feature such that the sum (of the same taxonomic level) is 1M and is done for obtaining more meaningful values for the LDA score. This package includes an internal Wilcoxon test and outputs discriminative features with abs LDA score > 2.0. Tests were performed using Treatment as class, for the three time points (P1, P2 and P3), and the taxonomic read counts considered subject vectors.

## Results

### Metagenome-based Metataxonomy

A metagenomic approach was used to investigate the effects of dietary supplementation with *Asparagopsis taxiformis* (ASP) on the dairy cattle microbiome, leading to phenotypic outcomes such as reduced CH_4_ emissions and altered volatile fatty acid (VFA) profiles. A total of 18 rumen samples were sequenced (in duplicate), representing two treatment groups (ASP-treated and control), with three animals per group sampled at three time points. Each sample was sequenced in technical duplicate, alongside positive and negative controls. The positive control showed the expected taxonomic profile consistent with the ZYMO mock community [Supplementary File 1], while the negative control yielded minimal reads numbering 428. All taxonomic and functional results are based on the number of reads mapped to metagenomically assembled contigs, or their coding regions.

On average, 82.93% of the contigs from each sample were annotated at the bacterial domain level, with a maximum of 88.11% and minimum 63.36% [Supplementary File 2]. When applying PCA analysis to the bacterial genus level taxonomy, no distinct clustering by treatment or time point was observed (Figure 1A). However, loose clustering by treatment emerged when 1 standard deviation ellipses were applied, and technical replicates consistently paired closely. PERMANOVA analysis indicated no significant overall differences in bacterial communities between treatments. However, significant differences were observed across time points, specifically between P3 vs. P1 and P3 vs. P2 (adjusted p = 0.003 for both comparisons; adjusted p > 0.05 for treatment effects) (Table 1), suggesting a time effect rather than a treatment effect.

**Figure 1.**
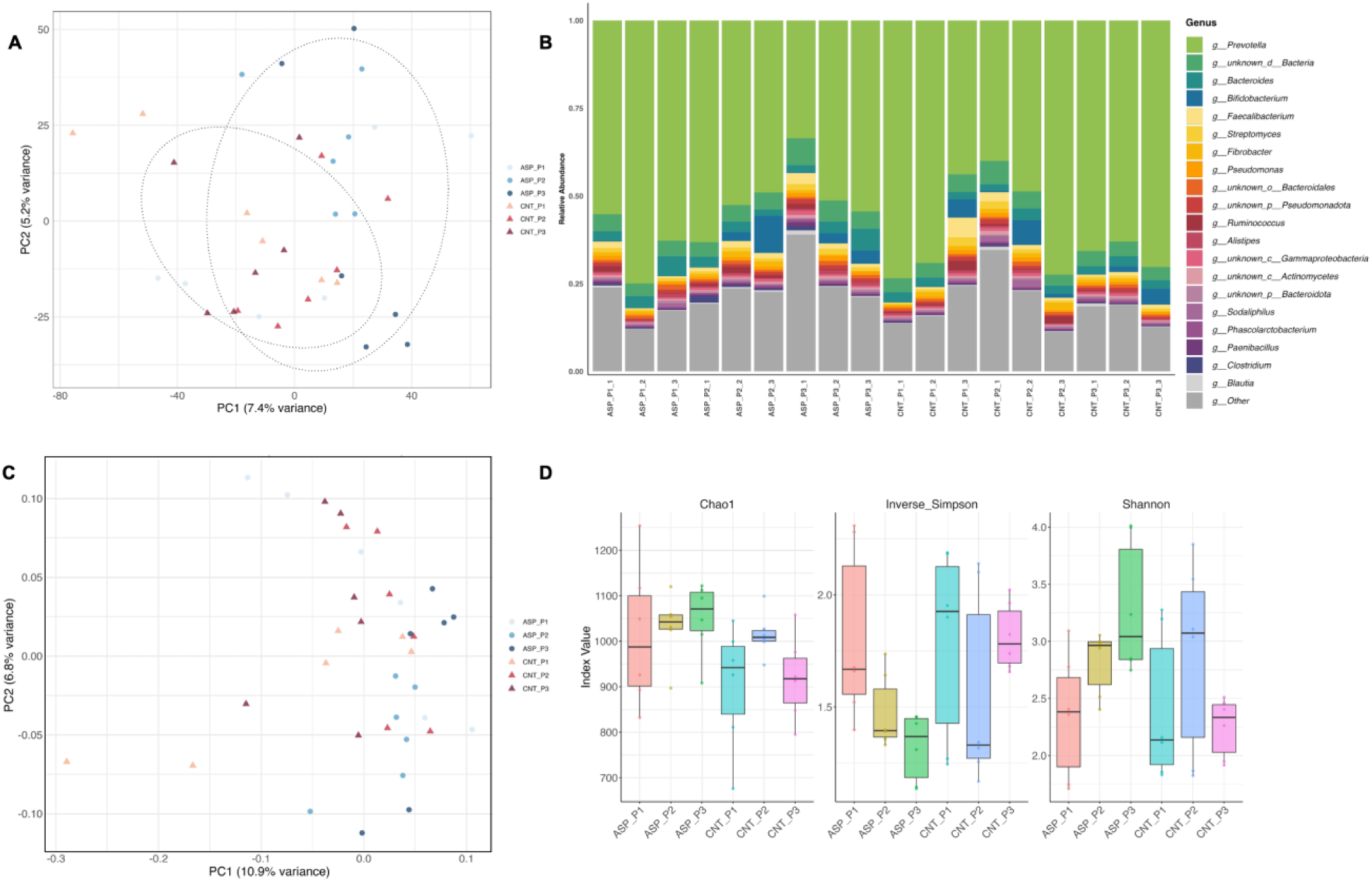
PCA of bacterial community composition at the genus level, based on metagenomic reads mapped to contigs from rumen fluid DNA. Samples are labelled by treatment (ASP, CNT), time point (P1, P2, P3) and technical replicated (1, 2), 1 SD ellipses were drawn around treatment groups. Figure 1B. Relative abundance of the top 20 bacterial genera across samples, based on metagenomic reads mapped to contigs with assigned taxonomy, all remaining genera are combined under “Other” for ease of visualisation. Figure 1C. Principal component analysis (PCA) of bacterial community composition based on Bray– Curtis dissimilarity. Analysis was performed at the genus level using relative abundance data from metagenomic reads mapped to contigs with assigned taxonomy. Samples are labelled by treatment, time point (e.g., ASP_P1, ASP_P2). Figure 1D. Alpha diversity metrics (Chao1, Inverse Simpson, and Shannon indices) calculated from genus-level bacterial taxonomic profiles based on metagenomic reads mapped to contigs. Box and violin plots display the distribution of diversity values across samples. Bars are coloured by treatment and time point (e.g., ASP_P1, ASP_P2) to visualise group-level trends.

**Table 1.**
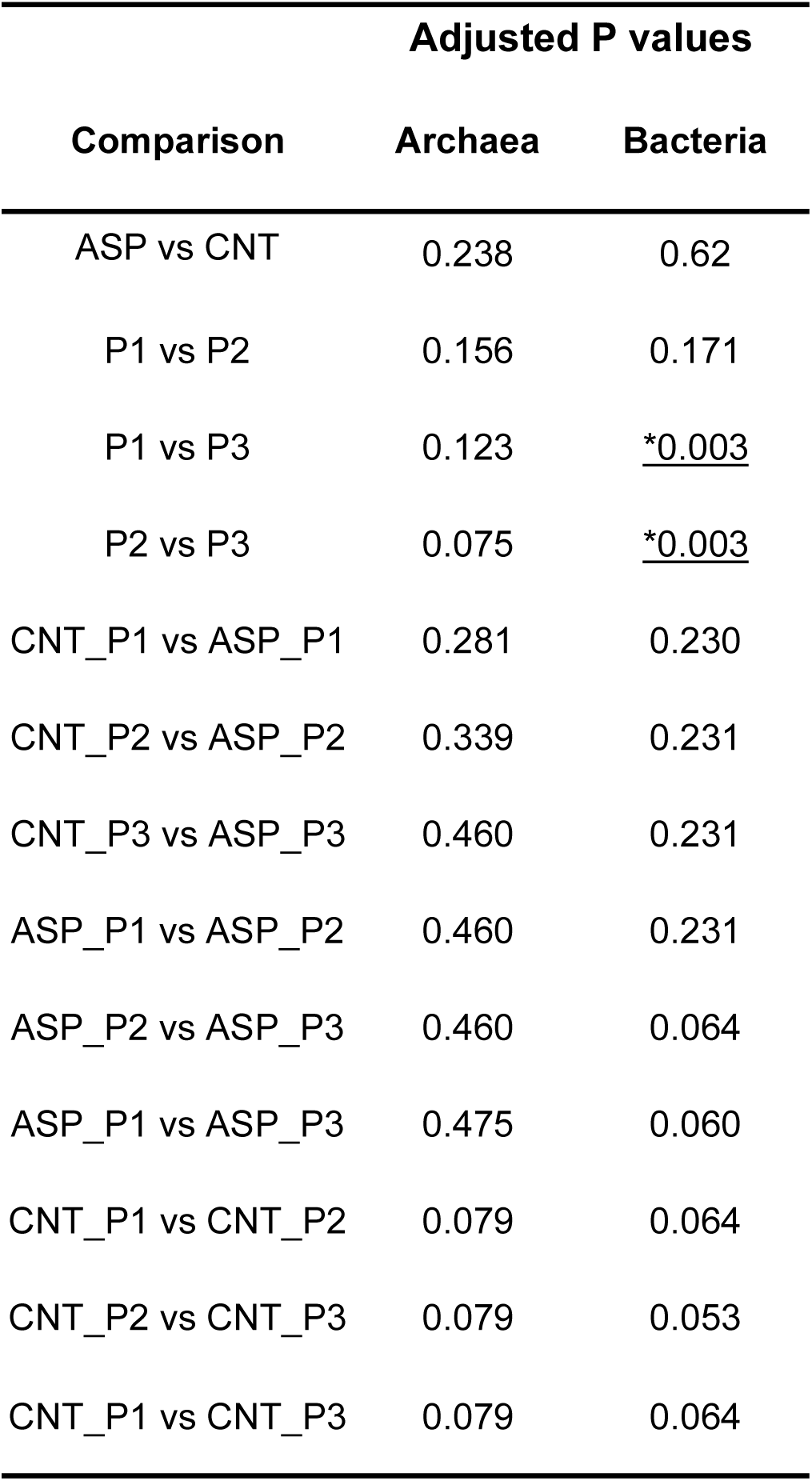
Statistical comparison of taxonomic and functional profiles based on metagenomic reads mapped to contigs, including bacterial and archaeal communities. P-values were calculated using PERMANOVA, adjusted using the Benjamini-Hochberg method. Significant p-values (p.adj >0.05) are underlined.

| Comparison | Adjusted P values |  |
| --- | --- | --- |
|  | Archaea | Bacteria |
| ASP vs CNT | 0.238 | 0.62 |
| P1 vs P2 | 0.156 | 0.171 |
| P1 vs P3 | 0.123 | <u>*0.003</u> |
| P2 vs P3 | 0.075 | <u>*0.003</u> |
| CNT_P1 vs ASP_P1 | 0.281 | 0.230 |
| CNT_P2 vs ASP_P2 | 0.339 | 0.231 |
| CNT_P3 vs ASP_P3 | 0.460 | 0.231 |
| ASP_P1 vs ASP_P2 | 0.460 | 0.231 |
| ASP_P2 vs ASP_P3 | 0.460 | 0.064 |
| ASP_P1 vs ASP_P3 | 0.475 | 0.060 |
| CNT_P1 vs CNT_P2 | 0.079 | 0.064 |
| CNT_P2 vs CNT_P3 | 0.079 | 0.053 |
| CNT_P1 vs CNT_P3 | 0.079 | 0.064 |

Considering phyla level, the most dominant phyla in all treatments and time points were Bacteroidota, Bacillota and Pseudomonadota with an average of relative abundance of 66.74%, 10.65%, and 9.23% respectively [Supplementary File 2]. The most abundant bacteria genus across all samples was *Prevotella* irrespective of time and treatment, with an average relative abundance of 0.58 (58.0%). Followed by *Bacteroides* (average 0.03, 2.98%), *Bifidobacterium* (average 0.02, 2.26%) and *Faecalibacterium* (0.015, 1.50%) (Figure 1B). Lefse analysis revealed two bacterial genera positively associated with control animals at P3 *Prevotella* and *Sphingobacterium*, whereas in ASP treated animals 59 genera were positively associated including *Oscillibacter, Alistipes, Lachnoclostridium, Enterocloster* and *Burkholderia* amongst others [Supplementary File 3]. Considering beta diversity based on the Bray-Curtis index, there was no clear clustering of the bacterial microbiome by time point or treatment (Figure 1C). Alpha diversity was evaluated using Chao1, inverse Simpson, and Shannon indices to capture different aspects of microbial community diversity. No significant differences were detected for Chao1 (p > 0.05), indicating similar species richness across groups and time points. When comparing CNT and ASP at P3, both inverse Simpson and Shannon indices showed significant differences (p.adj < 0.05) (Figure 1D), indicating that ASP treatment alters the structure and complexity of the microbial community, including richness and dominance.

At domain level there was an average of 137,517 reads assigned as archaea, with an average relative abundance of 1.35% of the data. The maximum reads per sample assigned to archaea was 753,711 (relative abundance of 4.84%), while the minimum per sample was only 6,401 (relative abundance of 0.06%). At the phylum level, taxonomic classification was high, with an average of only 0.003% relative abundance assigned as “unknown phylum” [Supplementary File 2]. The majority of archaeal sequences belonged to *Euryarchaeota*, with an average relative abundance of 95.75%. The next most abundant phylum was *Thermoproteota*, although its average relative abundance across samples was low (2.27%) [Supplementary File 2].

At the genus level, principal component analysis (PCA) showed that PC1 explained 8.6% of the variance. There was slight clustering of control samples on the right-hand side of PC1 (x-axis), with CNT_P3 clustering near the center. However, no distinct clustering was observed for ASP data when 95% confidence ellipses or 1 standard deviation ellipses were applied. Technical replicates did consistently pair together (Figure 2A). At the genus level, *Methanobrevibacter* was the dominant genus in most samples. By period 3, over 95% of the microbiome in control treatment samples was composed of *Methanobrevibacter* (Figure 2B).

**Figure 2.**
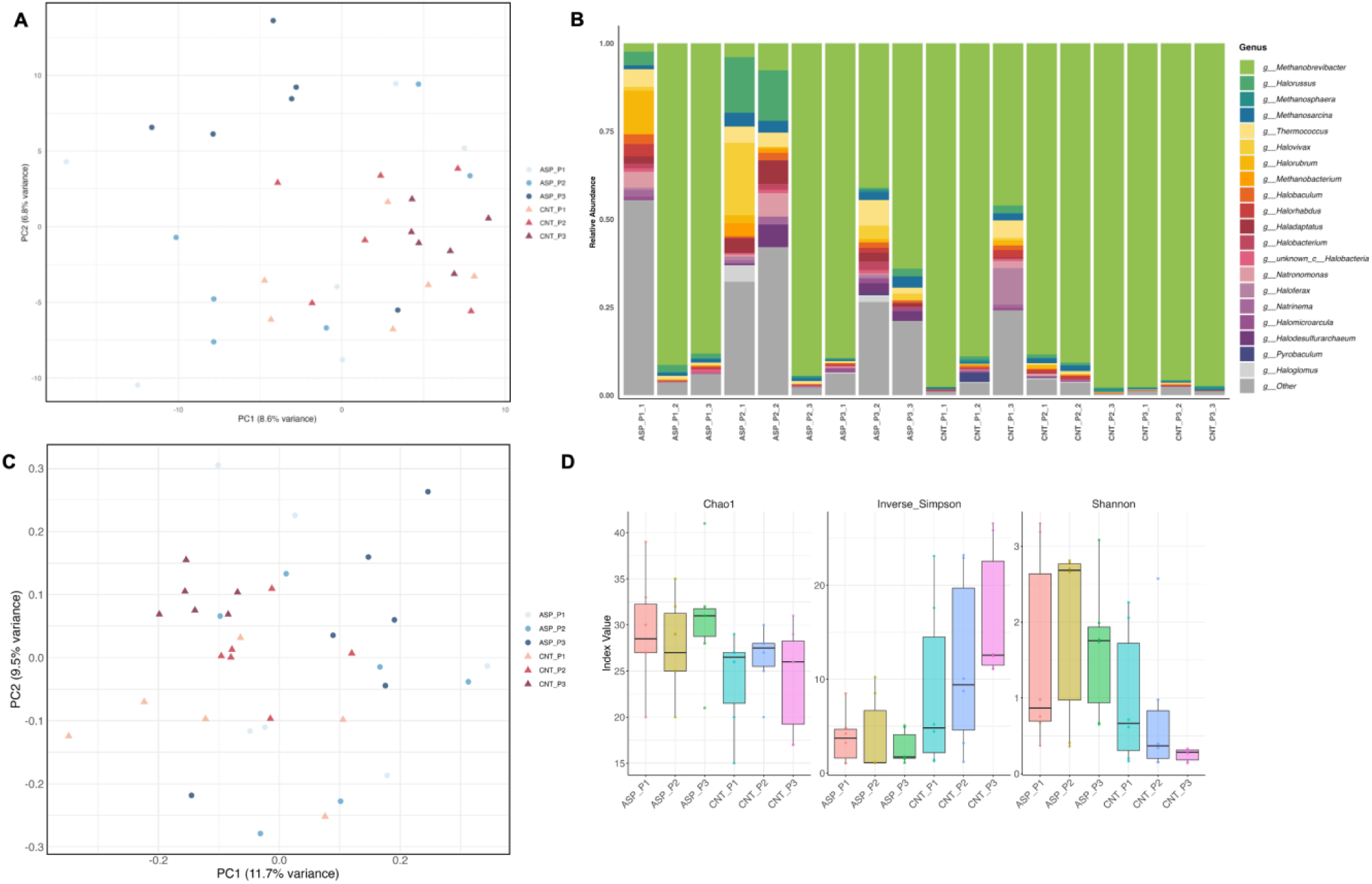
PCA of archaeal community composition at the genus level, based on metagenomic reads mapped to contigs from rumen fluid DNA. Samples are labeled by treatment (ASP, CNT), time point (P1, P2, P3) and technical replicated (1, 2), 1 SD ellipses were drawn around treatment groups. Figure 2B. Relative abundance of the top 20 archaeal genera across samples, based on metagenomic reads mapped to contigs with assigned taxonomy, all remaining genera are combined under “Other” for ease of visualisation. Figure 2C. Principal component analysis (PCA) of archaeal community composition based on Bray– Curtis dissimilarity. Analysis was performed at the genus level using relative abundance data from metagenomic reads mapped to contigs with assigned taxonomy. Samples are labeled by treatment, time point (e.g., ASP_P1, ASP_P2). Figure 2D. Alpha diversity metrics (Chao1, Inverse Simpson, and Shannon indices) calculated from genus-level archaeal taxonomic profiles based on metagenomic reads mapped to contigs. Box and violin plots display the distribution of diversity values across samples. Bars are colored by treatment and time point (e.g., ASP_P1, ASP_P2) to visualise group-level trends.

Similar clustering patterns were observed in the beta diversity analyses, with the only significant difference detected between CNT and ASP groups at P3 (p.adj = 0.007). This significant difference at P3 was also reflected in the alpha diversity metrics, with both Shannon and inverse Simpson indices showing significant differences between CNT and ASP (p.adj < 0.05). Over the course of the experiment, Shannon diversity significantly decreased over time in the control group, whereas this decline was not observed in the ASP-treated microbiomes. However, overall no significant differences were found at the archaeal genus level across comparisons, considering 95% confidence limits (p.adj > 0.05) (Table 1).

Exploring the archaeal community further, Linear Discriminant Analysis (LDA) revealed significant differences (Wilcoxon test and outputs discriminative features with abs LDA score > 2.0) between ASP and CNT treatments within each time point (Figure 3). At period 1 (P1), one archaeal genus was associated with each treatment: *Methanosphaera* was significantly associated with CNT, and *Acidianus* significantly associated with ASP, but neither were significantly linked to at treatment at either of the later periods (P2, P3). Focusing on P3, eight archaeal genera were significantly associated with the ASP treatment, including halophiles, Thermococcus (that was also significant at P2) and the methanogen *Candidatus Methanomethylophilus*. In contrast, only one archaeal genus, *Methanobrevibacter*, was significantly associated with the CNT treatment at P3.

**Figure 3.**
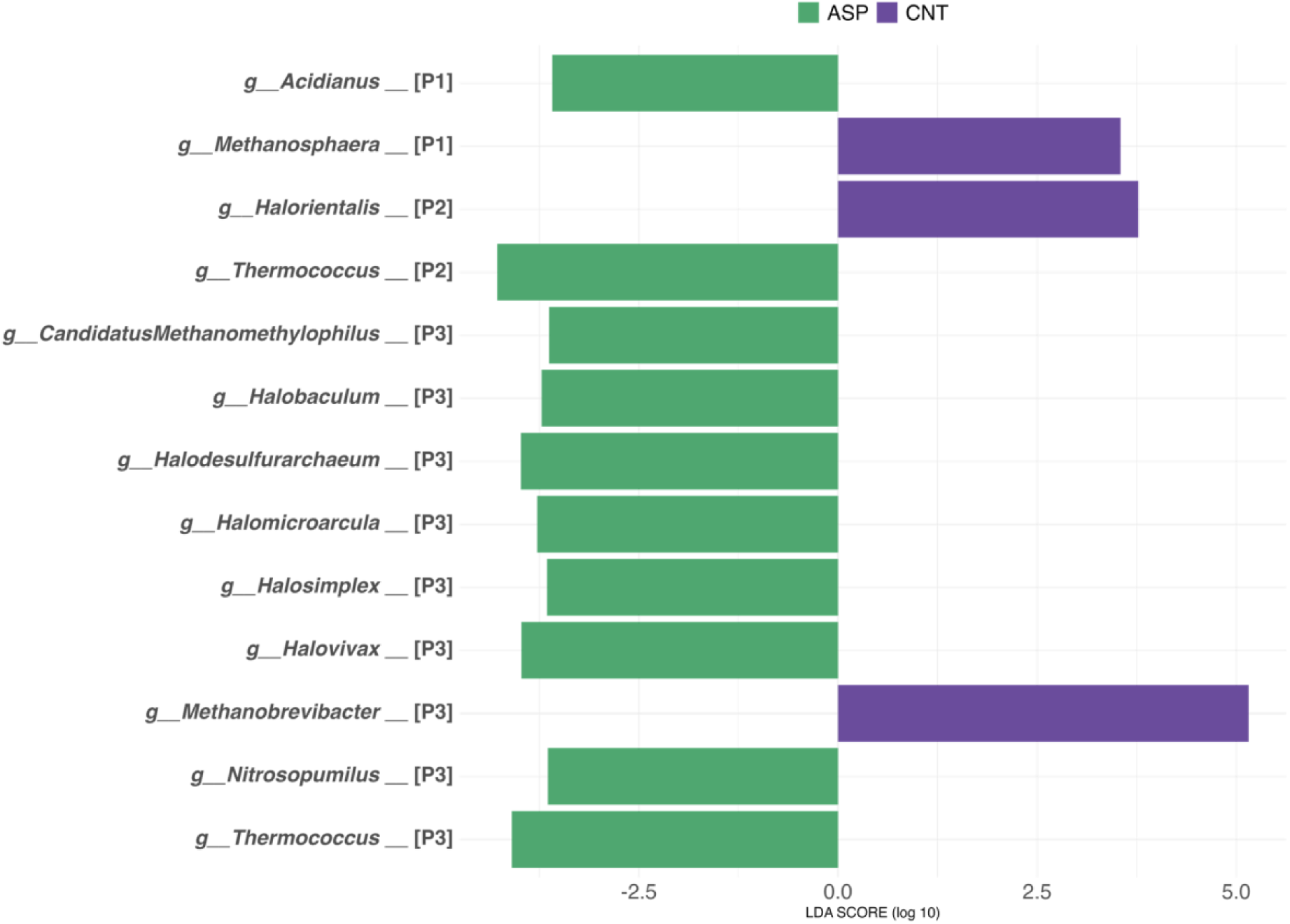
Differentially abundant archaeal genera identified using LEfSe (Linear discriminant analysis Effect Size), associated with ASP and CNT treatments across three time periods.. Only taxa with significant association with a timepoint are shown on the logarithmic LDA scores plot (Wilcoxon test and outputs discriminative features with abs LDA score > 2.0). Colors indicate the treatment group each archaeal genus is associated with based on LDA scores. Purple is associated with CNT animals, green is archaea associated with ASP treated animals.

To explore the methanogen population diversity further at a species level, in the metagenomes we employed a read mapping approach, using 27 methanogen genomes, and calculate the total mapped reads (to the methanogen) to the total number of reads per samples (MP:TR). We then investigated how the abundance of the methanogens changed over the course of the study.. This included 26 archaea species and 1 bacterial species *(Syntrophomonas wolfei* subsp. *wolfei* Goettingen) (Figure 4A). When comparing P3vsP1, the genomes with the largest differences over time were then explored further (Figure 4B).

**Figure 4:**
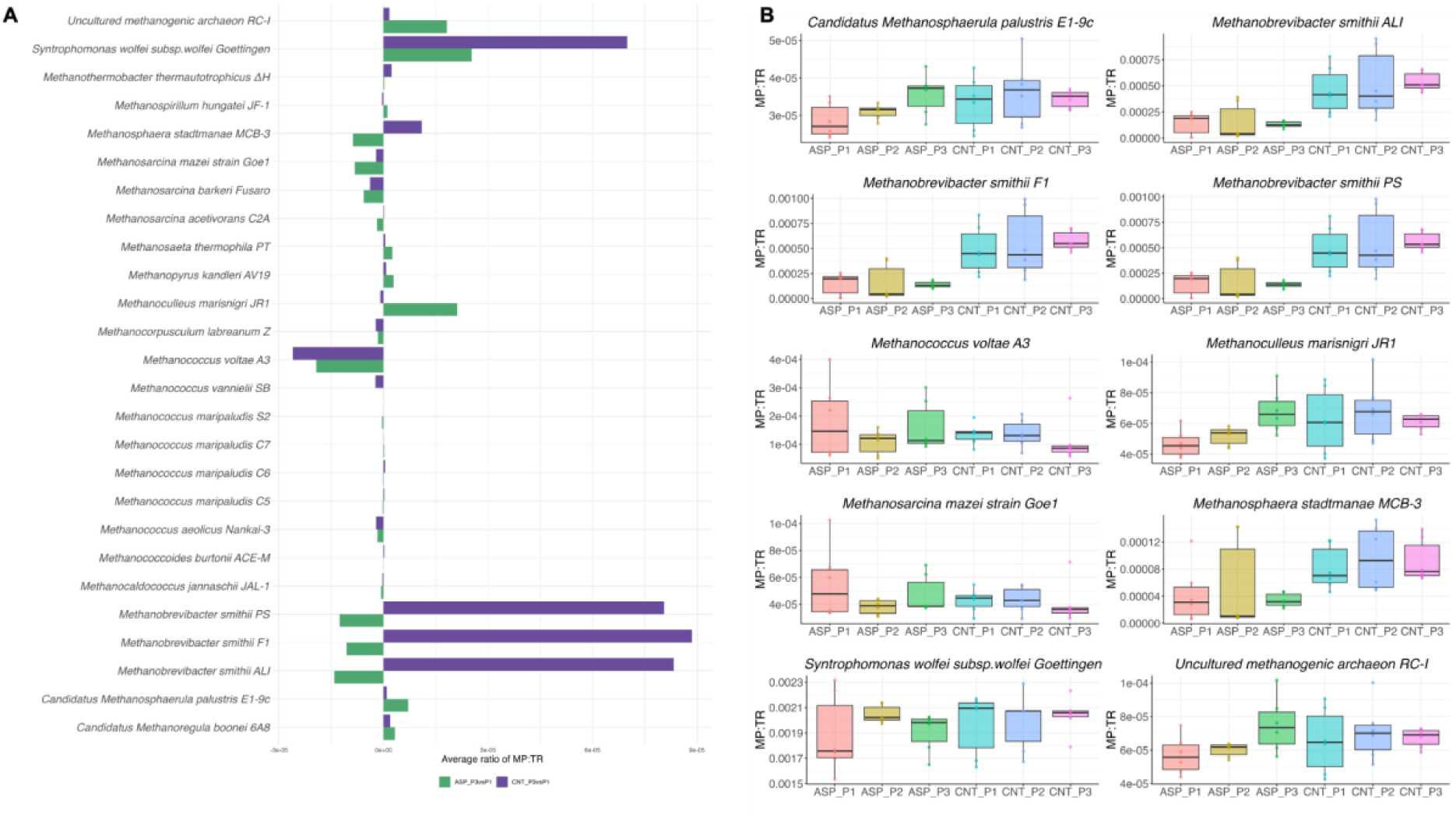
Changes in relative abundance of 27 reference genomes across treatments and time points, measured by the ratio of mapped reads to total reads (MP:TR). Metagenomic reads were mapped to each genome across two treatments and three time points. For each genome, mp:tr ratios were calculated per sample and averaged across animals. The bar chart displays the change in average MP:TR from time point 1 (P1) to time point 3 (P3), highlighting differences in genome abundance over time. Purple is associated with CNT animals, green is archaea associated with ASP treated animals. Figure 4B: Temporal dynamics of selected genomes with the largest changes in relative abundance. Genomes showing the greatest differences in MP:TR ratios between time point 1 (P1) and time point 3 (P3) were selected. For each genome, the MP:TR ratio (mapped reads to total reads per sample) is shown across all time points using box and violin plots. Each point represents an individual sample; boxes indicate the interquartile range, and violins show the distribution across replicates.

In ASP-treated animals, *Methanoculleus marisnigri* JR1 increased in relative abundance over time, Additionally, an uncultured methanogenic archaeon (RC-1) increased more substantially in ASP animals compared to CNT. Whereas *Methanobrevibacter smithii* (3 genomes) showed a modest decrease in ASP, as did *Methanosphaera stadtmanae* MCB-3 though this trend was variablE, particularly in ASP_P2 (Figure 4A and 4B). In CNT-treated animals, *Methanobrevibacter smithii* showed an overall increase in relative abundance over time. In contrast to ASP, *Methanosphaera stadtmanae* MCB-3 increased in CNT samples. The only bacterial genome included in the analysis, *Syntrophomonas wolfei* subsp. *wolfei* Goettingen, increased more markedly in CNT animals than in ASP, although its abundance was variable between individual animals (Figure 4A and 4B). *Finally, Methanococcus voltae* A3 decreased over time in both treatments at a similar rate.

### Metagenome-based functionality

Three functional categories were examined: KEGG Pathways, Enzyme Commission (EC) numbers, and CAZy (Carbohydrate-Active enZYmes) families [Supplementary File 2]. No significant differences were detected in KEGG Pathway profiles across treatments or time points (all p.adj > 0.05) (Table 2). In contrast, both EC profiles and CAZy enzyme distributions showed significant differences and were explored further.

**Table 2.**
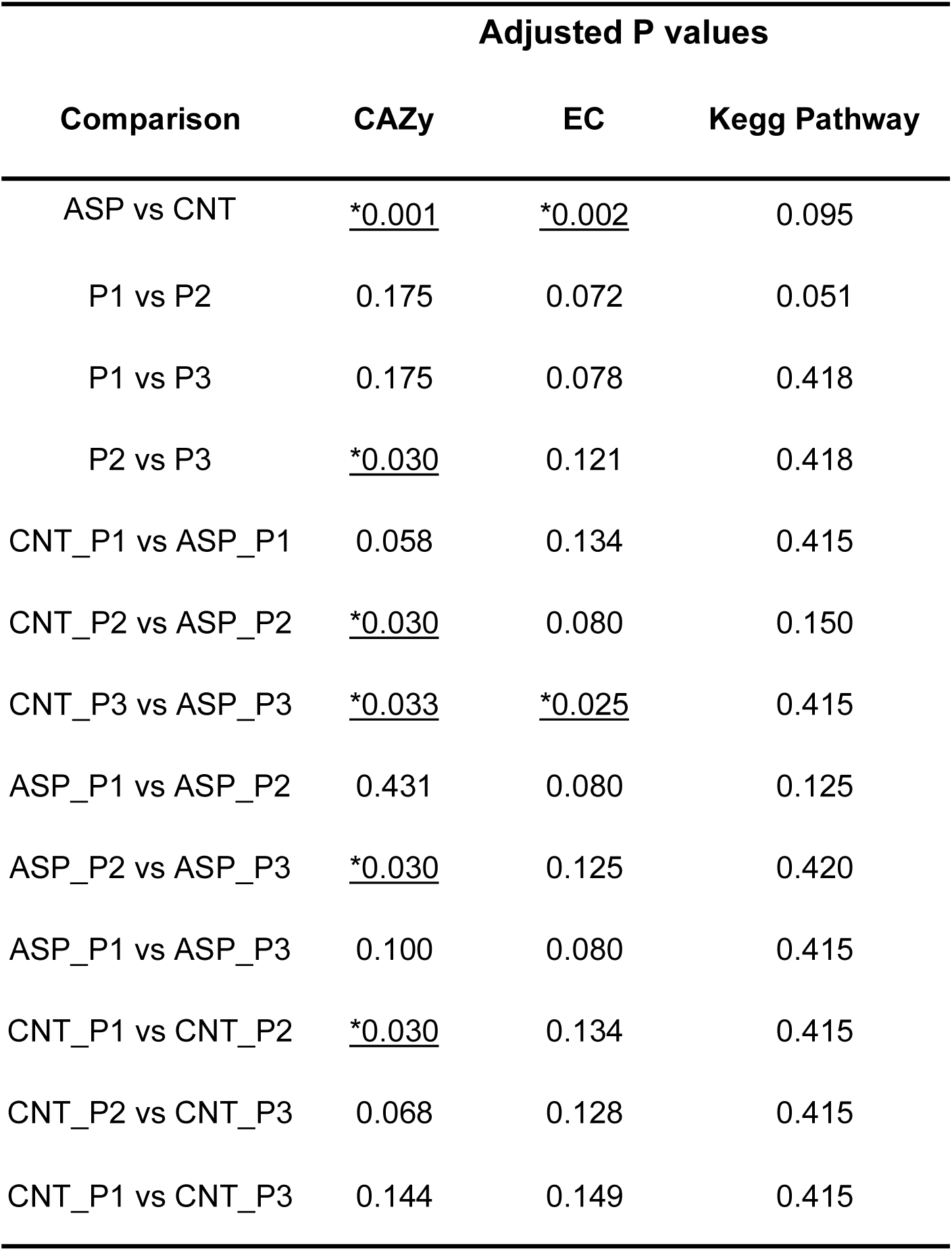
Statistical comparison of taxonomic and functional profiles based on metagenomic reads mapped to contigs, including bacterial and archaeal communities. P-values were calculated using PERMANOVA, adjusted using the Benjamini-Hochberg method. Significant p-values (p.adj >0.05) are underlined.

| Comparison | Adjusted P values |  |  |
| --- | --- | --- | --- |
|  | CAZy | EC | Kegg Pathway |
| ASP vs CNT | <u>*0.001</u> | <u>*0.002</u> | 0.095 |
| P1 vs P2 | 0.175 | 0.072 | 0.051 |
| P1 vs P3 | 0.175 | 0.078 | 0.418 |
| P2 vs P3 | <u>*0.030</u> | 0.121 | 0.418 |
| CNT_P1 vs ASP_P1 | 0.058 | 0.134 | 0.415 |
| CNT_P2 vs ASP_P2 | <u>*0.030</u> | 0.080 | 0.150 |
| CNT_P3 vs ASP_P3 | <u>*0.033</u> | <u>*0.025</u> | 0.415 |
| ASP_P1 vs ASP_P2 | 0.431 | 0.080 | 0.125 |
| ASP_P2 vs ASP_P3 | <u>*0.030</u> | 0.125 | 0.420 |
| ASP_P1 vs ASP_P3 | 0.100 | 0.080 | 0.415 |
| CNT_P1 vs CNT_P2 | <u>*0.030</u> | 0.134 | 0.415 |
| CNT_P2 vs CNT_P3 | 0.068 | 0.128 | 0.415 |
| CNT_P1 vs CNT_P3 | 0.144 | 0.149 | 0.415 |

For EC-level analysis, significant differences were observed overall between treatments (p.adj = 0.002), with specific differences detected at P3 when comparing ASP and CNT (p.adj = 0.025) (Table2). Principal Component Analysis (PCA) and Bray Curtis dissimilarity plots, used to quantify compositional differences between samples, revealed distinct separation between treatments: ASP samples were more dispersed along PC2, while CNT samples showed greater spread along PC1. Some clustering was observed within CNT_P3 samples, consistent with earlier observations.

Alpha diversity metrics for EC profiles also revealed treatment-specific trends. In ASP animals, the Shannon diversity index, an indicator of both richness (number of features) and evenness (distribution), increased substantially from P1 to P2 and remained elevated at P3. The inverse Simpson index, which emphasises dominant taxa and is more sensitive to abundance distribution, decreased over time in ASP animals. By P3, both Shannon and inverse Simpson indices were higher in ASP compared to CNT, suggesting greater functional diversity and a more even distribution of enzymatic capability [Supplementary File 4]. However, these differences were not statistically significant (p.adj > 0.05). The only significant alpha diversity change observed was within the CNT group, comparing P2 to P1 (p = 0.032). In contrast, beta diversity analyses revealed significant differences between P1 and P3 in the ASP group, as well as between ASP and CNT treatments at both P2 and P3 (p.adj < 0.05). Notably, the presence of significant differences in beta diversity but not alpha diversity indicates that while the overall enzymatic diversity within each sample remained relatively stable, the composition or relative abundance of enzymatic activities differed significantly between groups and over time. This suggests that ASP treatment influences the functional profile of the microbiome by altering enzyme composition rather than simply changing the total diversity of enzymatic functions.

Similarly, CAZy enzyme profiles showed significant overall differences between treatments (p.adj = 0.001). Significant differences were also observed at specific time points: P2 and P3 (p.adj = 0.030 and 0.033, respectively). Unlike EC, CAZy profiles also varied significantly over time. Differences were observed between P2 and P3 across all samples (p.adj = 0.030), and within-treatment comparisons showed significance for ASP between P2 and P3 (p.adj = 0.030) and for CNT between P1 and P2. However, this effect in CNT was not sustained from P2 to P3 (p.adj > 0.05) (Table 2).

Twenty-seven enzymes that were significantly different at P2 and P3, as identified by LEfSe analysis, and were further examined for their association with treatment and their potential interaction with methane metabolism. In control animals, several enzymes related to vitamin B12-dependent pathways were significantly more abundant (Figure 5A and 5B). These included enzymes involved in glycine and L-serine metabolism, as well as GTP conversion, which is linked to tetrahydromethanopterin (THMPT) and vitamin B12 coenzyme formation both with links methanogenesis. B12 is essential for enzymes like *methylmalonyl-CoA mutase* and *methionine synthase*, often linked to C1 metabolism and methyl group transfers. Enrichment of enzymes involved in glycine and L-serine metabolism suggests increased C1- unit generation, which feeds into folate and THMPT (tetrahydromethanopterin)-dependent pathways, key intermediates in methanogenic archaea.

**Figure 5.**
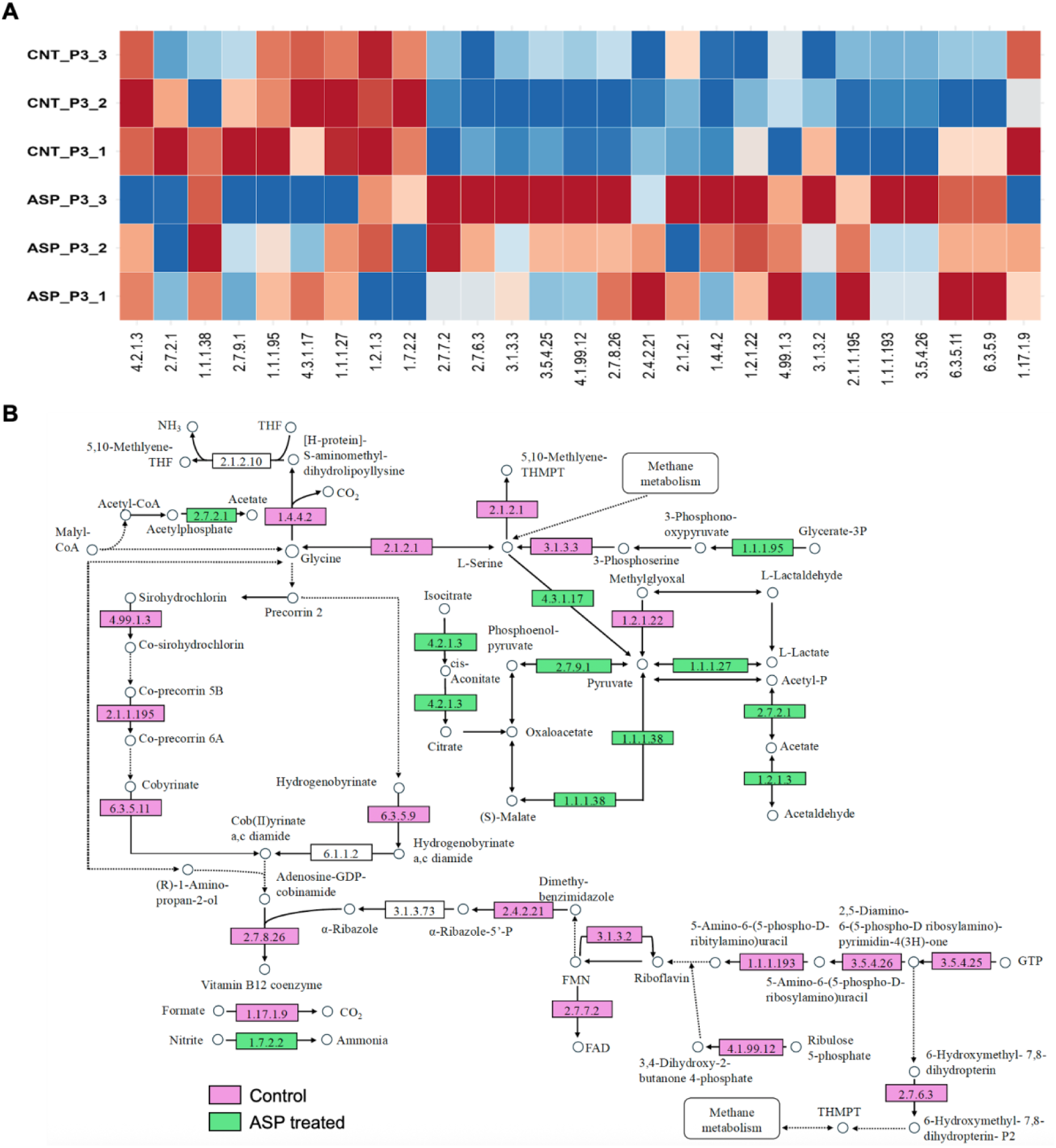
Number of metagenomic reads mapped to contigs annotated with EC numbers normalised to the total number of reads per sample, only those those involved in methanogenesis precursor pathways and were significantly different as determined by lefse are shown and those involved in methanogenesis precursor pathways. Full LDA scores for on KEGG Pathway, EC and CAZy can be found in Supplementary File 3. Here was focused on functions known to be associated with methane. Figure 5B. Pathways of EC with significant association (determined by lefse, pink CNT animals, Green ASP animals), pathways linked to methane metabolism.

The enzyme glycine dehydrogenase (EC 1.4.4.2), which mediates glycine cleavage to release CO₂, was notably elevated in control animals. CO₂ acts as the terminal electron acceptor in hydrogenotrophic methanogenesis, highlighting the enzyme’s potential role in supporting methane production. Additionally, the enzyme mediating formate conversion to CO₂ was also elevated (1.17.1.9), supporting increased one-carbon metabolism and methane-related pathways in the control group. It’s crucial in hydrogenotrophic methanogenesis, especially in anaerobic archaea and some bacteria, where formate serves as an electron donor for methane production. Formate dehydrogenases are also important in syntrophic bacteria that provide substrates (H₂, CO₂) for methanogens.

In contrast, ASP-treated animals showed increased abundance of the enzyme involved in acetyl-CoA conversion to acetate conversion (2.7.2.1), suggesting a shift in carbon flow away from methane-linked processes. Multiple enzymes in a number of pathways leading to pyruvate production were also more abundant in ASP-treated animals, including those converting L-lactate and isocitrate to pyruvate. Furthermore, elevated enzymes abundance able to produce pyruvate from acetate was observed, indicating a metabolic reprogramming toward energy production pathways less associated with methane synthesis.

There was no significant difference observed for any of the comparisons for the methyl-CoM reductase (MCR, EC 2.8.4.1), the enzyme that catalyses the final stage of methanogenesis.

By utilising MetaPont to extract and investigate the taxonomy of the contigs annotated with different EC numbers revealed a general trend, that in CNT animals, we observed a smaller number of taxa contributed to the functions shown in Figure 5, whereas in ASP-treated animals, a broader and more diverse range of taxa was involved. For example, among the four enzymes responsible for pyruvate production, ASP-treated animals exhibited reduced contribution from *Prevotella* and increased involvement of other genera such as *Bifidobacterium*, *Bacteroides*, and *Acidaminococcus*.

A similar pattern was observed in the enzymes associated with vitamin B12 coenzyme biosynthesis. In CNT animals, these functions were predominantly attributed to *Prevotella*, whereas in ASP animals, the activity was distributed across a wider set of taxa. For instance, enzyme EC 2.7.8.26 was largely attributed to *Prevotella* in CNT animals, but in ASP-treated animals, this function was instead carried out by *Solibacillus*, *Cubilibacter*, and *Bacillus*.

For enzyme EC 4.99.1.3, *Methanobrevibacter* was prominent in CNT animals but substantially reduced in ASP, with its role compensated by various other taxa including *Enterocloster* and *Pantoea*. A consistent pattern was observed with glycine metabolism-related enzymes. For EC 1.4.4.2 (glycine dehydrogenase), *Prevotella* was the main contributor in CNT animals, while in ASP animals, the function was more evenly shared among *Prevotella*, *Bacteroides*, and *Faecalibacterium*. Likewise, EC 2.1.2.1 (glycine hydroxymethyltransferase) showed *Prevotella*-dominance in CNT animals, but in ASP-treated animals, contributions came from *Prevotella*, *Marvinbryantia*, and *Bacteroides*. [Supplementary File 5].

## Discussion

A key goal for the ruminant sector is to develop innovative solutions to reduce GHG emissions thereby aiding planetary health and ensuring food security globally. Feeding ruminants *A. taxiformis* at levels up to 0.5% DMI have been shown to reduce CH_4_ emissions from ruminants by up to 80%, nonetheless the mechanism of action on the rumen microbiome is unclear. In order to develop this solution it is imperative that we understand the mechanism of action and ensure that there are on unintended consequences on an animals and a rumen microbiome level. While the exact mechanism of action to date has remained unclear, we propose that ASP exerts multifactorial effects, including enzyme inhibition, such as a reduction in vitamin B12 coenzyme biosynthesis, and altered activity of glycine dehydrogenase and other enzymes involved in one-carbon and nitrogen metabolism. Our results highlight that while the total functional potential related to the direct methanogenesis pathway may appear constant, less obvious pathways, particularly “feeder” pathways, that supply methanogenesis precursors were significantly altered, contributing to CH_4_ reduction in the animals.

The production parameters of this trial ((Krizsan et al., 2023), showed supplementation of ASP in dairy cattle resulted in lower feed intake but improved feed efficiency, along with a shift in fermentation products: reduced acetate and butyrate, increased propionate, lower CH_4_, and higher hydrogen production (Krizsan et al., 2023). In contrast, control animals showed higher feed intake, elevated acetate and butyrate production, higher CH_4_ emissions, and lower feed efficiency (Krizsan et al., 2023).

Supplementation with ASP has consistently been associated with reductions in CH_4_ emissions in ruminants, commonly attributed to the inhibition of *Methanobrevibacter* spp., dominant hydrogenotrophic methanogens that serve as key terminal electron acceptors in the rumen (Yergaliyev et al., 2024; Indugu et al., 2024). In agreement, in our study, we observed a marked reduction in the abundance of *Methanobrevibacter smithii* in ASP-treated animals, alongside a compensatory increase in potentially methylotrophic methanogens, particularly *Methanomethylophilus*. Although little is known about this uncultured genus, members of the *Methanomethylophilaceae* family have been associated with lower CH_4_ emissions and are known to utilise methylated compounds such as methanol and methylamines rather than hydrogen and CO₂ for methanogenesis (Krizsan et al., 2023; Poulsen et al., 2013).

Other studies have suggested ASP achieves CH_4_ reduction through inhibition of the MCR enzyme, similar to the mechanism of action of 3-Nitrooxypropanol (3-NOP) (Li et al., 2025, Indugu et al., 2025). However, our findings suggest that the mechanism underlying reduced CH_4_ emissions is multifactorial, involving both taxonomic changes and metabolic disruptions, particularly those related to vitamin B12 (cobalamin) and one-carbon metabolism. Vitamin B12 is an essential cofactor for several methanogenesis-related enzymes, including methyltransferases and methylmalonyl-CoA mutase, as well as enzymes involved in methionine synthesis and glycine cleavage (Raux et al., 2000; Warren et al., 2002). ASP has previously been hypothesised to interfere with B12-dependent processes, but our study is the first to identify specific enzymatic pathways potentially affected. This multifactorial mechanism of action is consistent with the relatively broad-spectrum antimicrobial activity of naturally occurring bromoform (Paul et al., 2006), considered one of the main active ingredients in red seaweed, especially when contrasted with the more targeted action of 3-NOP, which was specifically developed to inhibit the MCR enzyme. The CH_4_ reduction observed in this study (a 58% decrease in CH₄ intensity, Krizsan et al., 2023), despite no significant change in MCR enzyme abundance, was comparable to, or even exceeded, reductions reported in other studies investigating ASP supplementation (an average 39.0% reduction in CH₄ intensity; Kebreab et al., 2025)

In ASP-treated animals, we observed a reduction in the abundance of vitamin B12 coenzyme biosynthesis-related enzymes and a diversification of microbial taxa contributing to these pathways. While control animals exhibited B12-associated functions predominantly carried out by *Prevotella*, ASP-treated animals showed a more distributed taxonomic contribution involving *Solibacillus*, *Cubilibacter*, and *Bacillus*. This suggests a possible microbial compensation for the disruption of B12-dependent metabolism, potentially due to direct or indirect inhibition of B12 synthesis by ASP or its bioactive compounds. Previous studies have shown that high ruminal concentrations of vitamin B12 are associated with increased abundance of *Prevotella* and *Methanobrevibacter i*n the rumen and throughout the gastrointestinal tract of ruminant animals (Franco-Lopez et al., 2020; Jiang et al., 2022). Whereas low B12 levels correlate with higher relative abundances of *Bacteroidetes*, *Ruminiclostridium*, *Butyrivibrio*, and *Succinimonas (*Franco-Lopez et al., 2020). These findings support our interpretation that ASP treatment disrupts vitamin B12 biosynthesis and drives a compensatory shift in the taxonomic contributors to this essential metabolic function. This diversification of taxa contributing to vitamin B12 biosynthesis in ASP-treated animals indicates niche displacement, where alternative microbes assume metabolic roles previously dominated by a narrower group.

This observed functional redistribution reflects both microbial resilience and redundancy within the rumen microbiome, enabling essential processes to persist despite targeted disruption. Genes within the microbiome do not strictly belong to fixed taxa; instead, they can be transferred between organisms, shifting their functional contributions in response to selective forces. Thus, microbiome resilience must be considered not only at the level of genomes or taxa, but also at the level of individual genes. Although metagenomic assembly tools such as the one used in this study, Metaspades, uses k-mer abundance and read coverage to try and ensure that each contig is derived from a single ‘species’, they are not always accurate. Therefore, it is plausible that certain genes that are more present in a sample than its neighbors on the same contig will have more reads mapped to it and thus provide important information on the abundance of individual genes. Such functional plasticity underscores the adaptive capacity of the microbiome and may explain why efforts to reduce methane emissions through microbiome-targeted interventions can underperform.

The potential mechanism of action mirrors that of known halogenated CH_4_ analogues, such as bromoform, a major active component of ASP. These compounds have been shown to irreversibly react with reduced vitamin B12, effectively inhibiting cobamide-dependent methyltransferases that are central to methanogenesis (Czerkawski & Breckenridge, 1975; Demeyer & Van Nevel, 1975; Wood et al., 1968). Such inhibition would disrupt the final methyl-transfer reactions in methanogenic archaea (Indugu et al., 2024; Kanno et al., 2022). Moreover, bromoform and related compounds may also impair the synthesis of methionine from homocysteine, a process likewise dependent on cobamide cofactors (Brot & Weissbach, 1965; Wood & Wolfe, 1966).

The observed decrease in glycine dehydrogenase (EC 1.4.4.2) activity in ASP-treated animals further supports a disruption in one-carbon metabolism. In control animals, elevated glycine cleavage likely contributed to increased CO₂ availability, feeding hydrogenotrophic methanogenesis. The reduced activity of this pathway in treated animals may contribute to lowered CH_4_ by limiting substrate availability. This is consistent with broader metabolic shifts observed in low methane-emitting ruminants, where differences in carbohydrate metabolism and microbial defence pathways have also been reported. For example, sheep with divergent CH_4_ emission phenotypes showed differential abundance of enzymes involved in glucose and lactate metabolism, including components of the methylglyoxal pathway (a glycolytic bypass that results in D-lactate production) (Bond et al., 2022).

Similarly, formate dehydrogenase (EC 1.17.1.9), another CO₂-generating enzyme supporting methanogenesis, was diminished under ASP treatment, indicating a broader suppression of formate- and glycine-derived C1 metabolism. This finding aligns with prior observations that hydrogen (H₂) and formate serve as critical intermediates in anaerobic ecosystems, including the rumen, where they function as electron sinks for primary fermenters in the absence of external electron acceptors (Kelly et al., 2022). The accumulation or depletion of these intermediates can profoundly affect fermentative balance and microbial growth. In particular, methanogens play a key role in consuming both H₂ and formate to maintain the redox equilibrium necessary for efficient fermentation. Experimental evidence from environments such as sewage digesters, biogas reactors, and acidic peatlands suggests that hydrogen and formate pools tend to be rapidly equilibrated and energetically interconvertible, reflecting a high degree of microbial regulation (Schink et al., 2017; Hunger et al., 2016). In ruminants, methanogens have been shown to utilize formate alongside hydrogen and other substrates, although *in situ* metatranscriptomic data suggest that formate-dependent H₂ production is not a major pathway under normal conditions (Greening et al., 2019). Nevertheless, co-culture studies demonstrate that formate can serve as a supplementary electron donor, especially when transferred from fermentative bacteria like *Ruminococcus albus* (Greening et al., 2019). Furthermore, methanogens such as *Methanococcus maripaludis* contain multiple formate dehydrogenase genes that redundantly support growth on formate, underscoring its central role in methanogenic metabolism (Wood et al., 2003). The observed suppression of formate dehydrogenase expression under ASP treatment may therefore represent a targeted disruption of interspecies electron transfer pathways, particularly those facilitating methanogenesis, and could contribute to the broader inhibition of ruminal hydrogen and formate fluxes.

Additionally, our data suggest metabolic rerouting in ASP-treated animals. A number of enzymes involved in acetate conversion to pyruvate, including acetate kinase (EC 2.7.2.1), were more abundant, potentially reflecting a shift in carbon flux away from acetate accumulation and CH_4_ generation and toward alternative energy-yielding pathways. Given that pyruvate is a key precursor for propionate synthesis, this shift may help explain the increased propionate and reduced acetate and CH_4_ concentrations observed in previous ASP feeding studies (Krizsan et al., 2023). This is supported by findings from low methane-emitting ruminants, where microbial communities show lower abundances of acetate-forming enzymes such as acetate kinase and phosphotransacetylase, and altered pyruvate metabolism routes that reduce acetyl-CoA and acetate production in favor of alternative pathways (Wallace et al., 2015). Methanogenesis inhibition is also known to increase propionate formation via hydrogen-consuming pathways, thereby mitigating hydrogen accumulation, which can otherwise impair rumen fermentation by disrupting NADH/NAD⁺ recycling (Romero et al., 2023; Camer-Pesci et al., 2023). These regulatory shifts in carbon and hydrogen flux may reflect a broader microbial adaptation to maintain redox balance and fermentation efficiency under ASP-induced suppression of methanogenesis.

## Conclusion

Overall, our results indicate that ASP reduces methane production not only through direct methanogen inhibition but also by targeting cobamide-dependent metabolic networks and shifting rumen fermentation dynamics. For the first time, we predict the specific enzymes, involved in vitamin B12 (cobamide) coenzyme biosynthesis, that are suppressed by ASP treatment. This includes disruption of B12-dependent and C1 metabolic pathways, alongside a reprogramming of carbon flow toward pyruvate and propionate rather than acetate and methane. We also provide novel taxonomic resolution of the microbial contributors to these functional shifts, linking enzymatic activity to microbial taxa, a level of insight rarely achieved outside of metagenome-assembled genome (MAG)-based studies. By employing a holistic approach that integrates taxonomic, functional, and niche displacement analyses, we offer a more comprehensive overview of the microbiome changes induced by *Asparagopsis* supplementation in ruminants. These findings advance a more mechanistic understanding of how red seaweed supplementation influences microbial ecology and metabolic function in the rumen, with important implications for sustainable ruminant production.

## Supplementary Material

Supplementary File 1 - Taxonomy of the positive control sample

Supplementary File 2 - Raw reads- full lineage taxonomy, domain, KEGG pathway, EC, CAZy

Supplementary File 3 - Lefse results- bacteria, archaea, KEGG pathway, EC, CAZy

Supplementary File 4 - Function analysis figure panel: EC A- PCA, B- Bray Curtis, C- Alpha diversity. CAZy D- PCA, E- Bray Curtis, F- Alpha diversity

Supplementary File 5 – Taxonomic data assigned by Kraken2 of the contigs annotated with enzymes of interest, data was mined using MetaPont

---

**Supplementary File 4:**
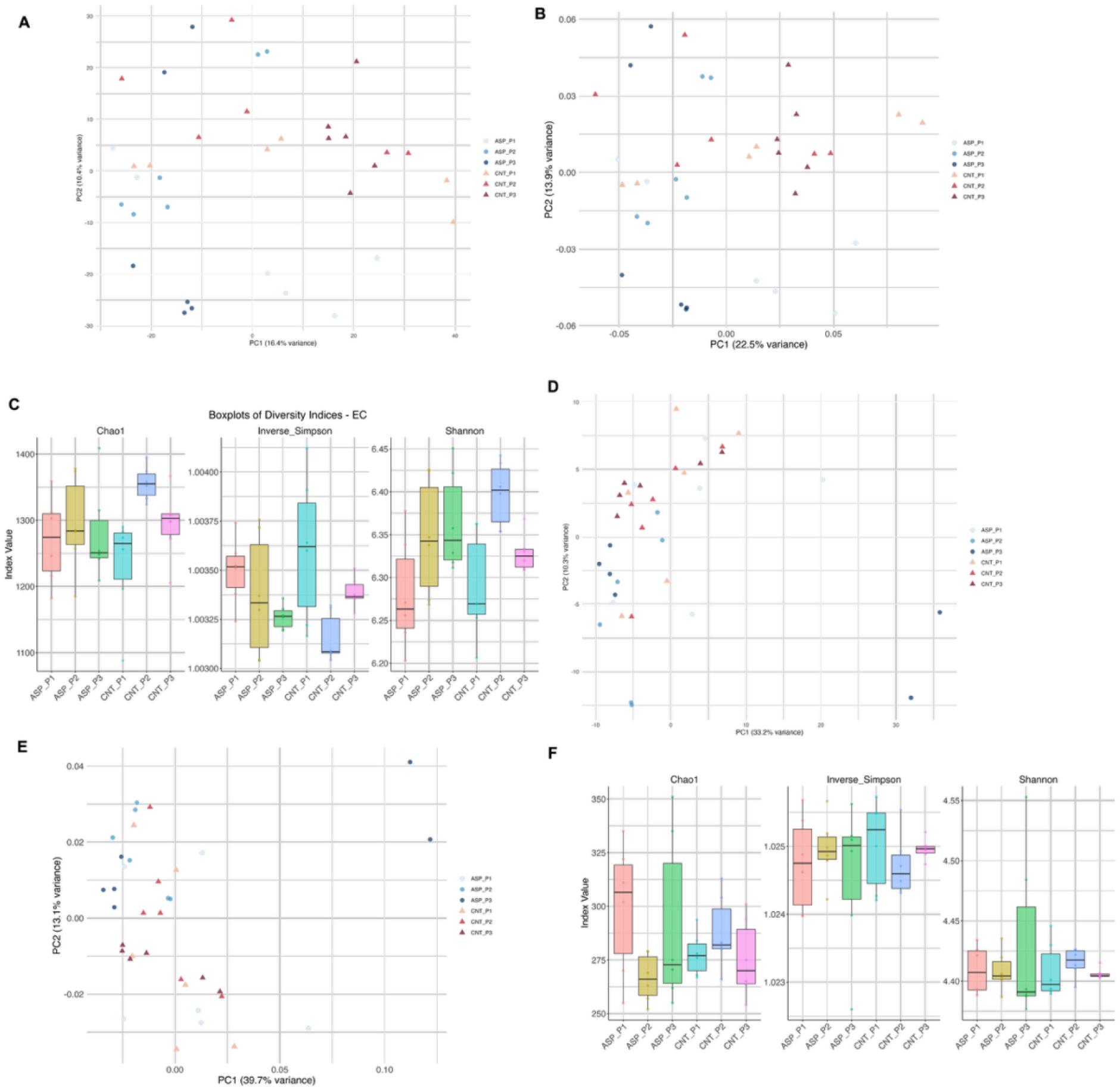
Function analysis figure panel: EC A- PCA, B- Bray Curtis, C- Alpha diversity. CAZy D- PCA, E- Bray Curtis, F- Alpha diversity

